# A fusion Cell-Permeable C16orf74 Peptide Selectively Disrupts Calcineurin-NFAT Interaction and Inhibits T-cell Activation Without Cytotoxicity

**DOI:** 10.64898/2026.05.21.726749

**Authors:** Adi Cohen, Matan Gabay, Sumit Gupta, Marina Sova, Daniel Bar, Jerome Tubiana, Maayan Gal

## Abstract

Calcineurin (Cn) is a protein phosphatase that initiates T-cell activation by dephosphorylating the transcription factor NFAT, driving its nuclear translocation and the transcription of immune-related genes. While clinical immunosuppressants like Cyclosporine A (CsA) potently inhibit Cn, they completely block its catalytic site, leading to non-specific inhibition and severe off-target toxicity. Selectively targeting the specific protein-protein interaction (PPI) between Cn and NFAT presents a safer therapeutic strategy. We previously identified the C16orf74 (C16) peptide as a high-affinity Cn-NFAT PPI inhibitor; however, its utility in cellular systems is restricted by poor membrane permeability. In this study, we evaluated cell-penetrating peptide (CPP) conjugates of C16 with an N-terminus transactivator of transcription (TAT) and polyarginine (R11) to enable efficient intracellular delivery. Structural modeling, fluorescence polarization displacement, and pull-down assays confirmed that the CPP–C16 conjugates retain the ability to compete with an NFAT-derived peptide and bind Cn. Fluorescence microscopy demonstrated efficient intracellular entry of TAT-C16 and R11-C16 in mammalian cells, and effective inhibition of NFAT nuclear translocation and attenuation of downstream NFAT-dependent transcriptional activity of the IL-2 gene in human T cells at concentrations of 10 µM or lower. Crucially, unlike CsA, the CPP-C16 peptides exhibited minimal cytotoxicity even at high concentrations of up to 50 µM, establishing a potential safe therapeutic window. These findings establish CPP-C16 conjugates as effective, cell-permeable, and non-toxic inhibitors of the Cn-NFAT signaling axis, providing the basis for the development of PPI-directed immunosuppressants.

## Introduction

Calcineurin (Cn) is a Ca^2+^ calmodulin-dependent serine–threonine phosphatase that regulates T-cell activation [1–3]. Following T-cell receptor engagement, a rise in intracellular calcium levels triggers Cn activation, which in turn dephosphorylates the transcription factor Nuclear Factor of Activated T-cells (NFAT), driving its translocation to the nucleus. There, NFAT induces the transcription of essential immune cytokines, such as interleukin-2 (IL-2), to promote T-cell proliferation and immune response [4–6]. Cn is a heterodimer composed of a catalytic A (CnA) and a regulatory B (CnB) subunit (Fig. 1A) [7,8]. While Cn dephosphorylates a diverse array of protein substrates [9–13], NFAT is recognized among its primary physiological targets. Efficient dephosphorylation requires NFAT to dock onto Cn via two distinct epitopes, both located distally from the catalytic center (red surface, Fig. 1A). The first interaction involves the NFAT PxIxIT motif (Fig 1B. CN-1) binding to CnA (green surface, Fig. 1A), while the second involves the NFAT LxVP motif (Fig. 1B, CN-2) binding at the CnA–CnB interface (blue surface, Fig. 1A) and can also compete with the PxIxIT motif [14–16].

**Figure 1.**
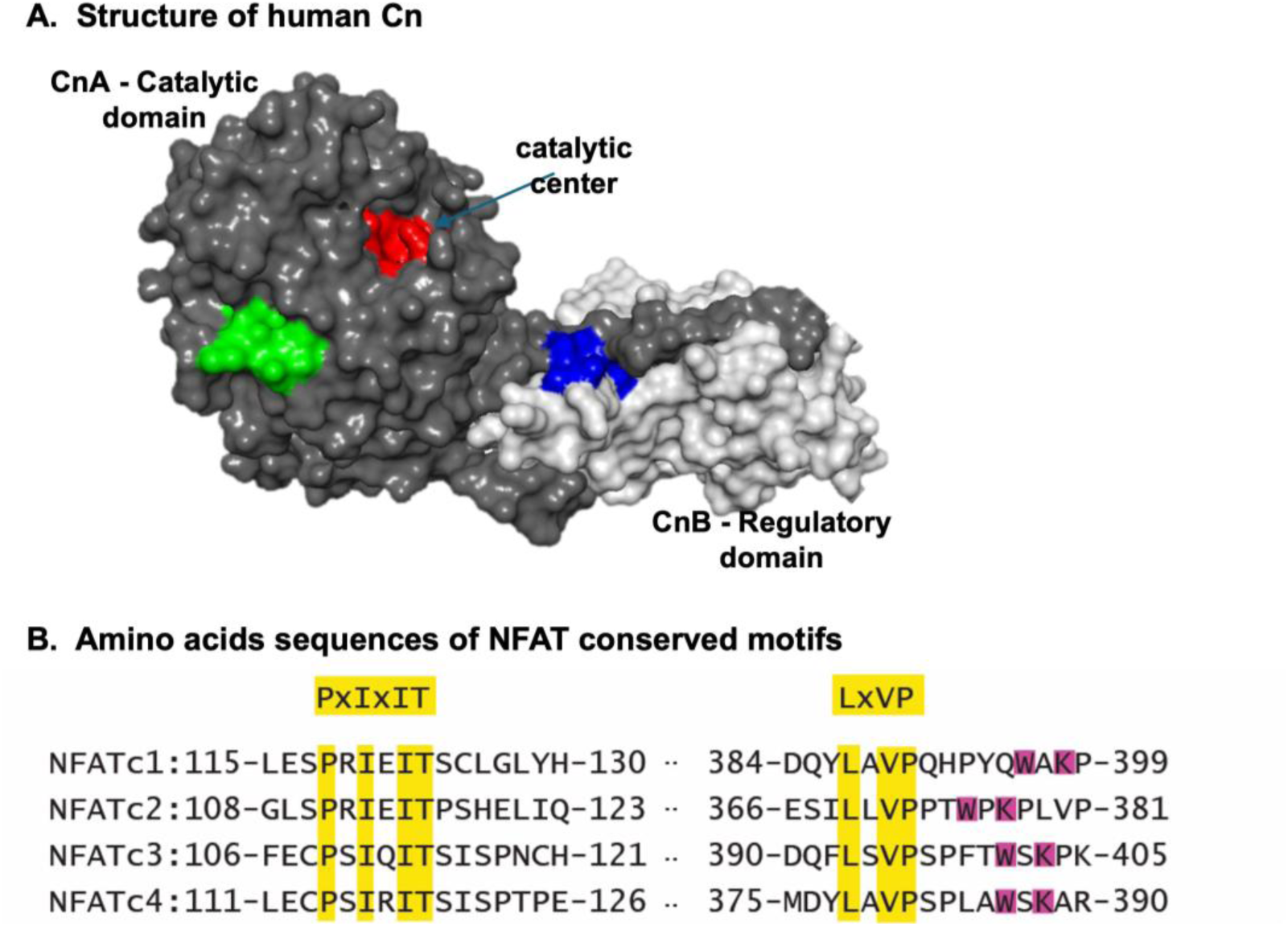
Structural organization and functional motifs of the Cn–NFAT interaction. **(A)** Surface representation of the Cn heterodimer, comprising the catalytic A subunit (CnA, dark gray) and the regulatory B subunit (CnB, light gray). Key functional regions are highlighted: the catalytic center (red); the PxIxIT-binding epitope on CnA (green); and the LxVP-binding interface at the CnA–CnB junction (blue). **(B)** Sequence alignments of NFAT isoforms display the primary PxIxIT and LxVP docking sequences, where ’x’ denotes variable amino acids. The functional WxK motif, located within the LxVP region, is also highlighted. The PxIxIT and LxVP segments flank NFAT phosphorylation sites for efficient dephosphorylation by Cn.

The central role of Cn–NFAT signaling in T-cell function marks it as a primary target for immunosuppressive therapy. Indeed, Cn inhibitors such as Cyclosporine A (CsA) and Tacrolimus (FK506) are widely used in the clinic [17,18]. While highly potent, the inhibition of Cn’s catalytic site indiscriminately blocks the phosphatase activity against all cellular substrates, leading to severe off-target effects, most notably nephrotoxicity and neurotoxicity [19,20]. Consequently, there is a need for molecular strategies that selectively disrupt NFAT dephosphorylation by Cn while potentially enabling Cn’s activity against other substrates.

An alternative approach is the targeted inhibition of the Cn–NFAT protein-protein interaction (PPI) [21–23]. Because this interface relies on specific docking surfaces distinct from the active site, it represents a highly validated, druggable target. Exploiting this structural feature led to the development of the ‘PVIVIT’ peptide, a high-affinity optimized variant of the native NFAT PxIxIT motif that selectively inhibits NFAT-driven transcription [24,25]. Cell-permeable derivatives have since demonstrated immunosuppressive efficacy in models of autoimmunity and cancer [26–28].

Despite significant progress in developing Cn-NFAT peptide inhibitors, the majority of high-affinity candidates rely on the canonical, NFAT-derived PVIVIT sequence. To expand this therapeutic space, our previous work utilized an integrative computational design and experimental screening approach to identify novel Cn-NFAT inhibitors derived from alternative substrates. This strategy culminated in the discovery of the C16orf74-derived peptide (C16) as a potent inhibitor [29]. Recent studies have characterized C16orf74 protein (also named Calcimembrin) as a unique physiological regulator of Cn. It utilizes an intrinsically disordered region to anchor the phosphatase to intracellular membranes via a distinct mode of interaction [30]. This structural basis further supports the potential of the C16 peptide not only as a membrane-targeting anchor but also as a basis for designing high-affinity inhibitors that potentially simultaneously exploit multiple docking surfaces on Cn, making it a promising inhibitor of NFAT activation. However, as the plasma membrane is impermeable to most unmodified peptides, the utility of C16 in cellular systems is fundamentally limited by its delivery into the cell. To overcome this barrier we conjugated the C16 peptide to the well-established cell-penetrating peptides (CPPs), the HIV-1 Trans-Activator of Transcription (TAT) and the polyarginine cation (R11) peptides [31,32]. The latter are part of the CPP family with a short cationic sequence shown to enable rapid and efficient cellular internalization of biomolecules. In this study, we evaluate the cellular uptake, binding and intracellular functional efficacy of these fusion peptides as targeted inhibitors of the Cn-NFAT signaling axis.

## Materials and Methods

### Peptides

All peptides were synthesized with C-terminal amidation by GL Biochem (Shanghai, China). Peptides were dissolved in DMSO and diluted to working concentrations before use. Peptide sequences were the following: (C16) KHLDVPDIIITPPTPT; (TAT-C16) GRKKRRQRRRPQKHLDVPDIIITPPTPT and (R11-C16) RRRRRRRRRRRKHLDVPDIIITPPTPT.

### GST-fusion peptides expression and purification

GST, GST-PVIVIT and GST-C16 fusion proteins were expressed in *Escherichia coli* BL21 cells as described earlier [15]. Briefly, bacterial cultures were grown to an OD600 of approximately 0.8 and protein expression was induced with 1 mM IPTG overnight at 25°C. Bacterial pellets were lysed in PBS containing protease inhibitor (Merck, 11836170001), and GST-tagged proteins were purified using an AKTA purification system with a GSTrap HP column (Cytiva). Purified proteins were dialyzed against PBS, centrifuged at 18,000 × g for 20 min to remove aggregates, and protein concentration was determined using NanoDrop spectrophotometry and BCA assay (Thermo fisher, 23225).

### GST pull-down assay

For pull-down experiments, 8 × 10^7^ Jurkat E6-1 cells were collected, washed with PBS, and lysed in RIPA buffer containing 1:100 protease inhibitor. For each condition, 30 µL Glutathione Sepharose 4B beads (Cytiva, 17-0756-01) were washed and incubated with 25 µg GST-tagged proteins or beads only as a negative control for 2 h at 4°C on a rotator. After removal of unbound proteins, Jurkat cell lysates were added to the beads and incubated for 3 h at 4°C on a rotator. Beads were then washed with RIPA buffer containing protease inhibitor, and bound proteins were eluted using 4× sample buffer followed by boiling at 95°C for 10 min prior to western blot analysis.

### Western blotting

Proteins were separated by SDS-PAGE using 12% acrylamide gels (1.5 mm thickness) and transferred onto PVDF membranes (Bio-Rad, 1704157). Membranes were then probed for detection of GST-tagged proteins using a mouse anti-GST antibody (GenScript, A00865) and for CnA using a rabbit anti-CnA antibody (Abcam, ab52761). HRP-conjugated secondary antibodies were used for detection: goat anti-mouse HRP (Sigma-Aldrich, AP130P) for mouse primary antibody and goat anti-rabbit HRP (Sigma-Aldrich, 12-348) for rabbit primary antibody.

### Cells and culture conditions

HeLa cells (ATCC, CCL-2) and Jurkat E6-1 cells (ATCC, TIB-152) were used in this study. Cells were maintained at 37°C in a humidified incubator with 5% CO₂. HeLa cells were cultured in high-glucose DMEM (Sartorius, 01-052-1A) supplemented with 10% fetal bovine serum (FBS, Diagnovum, D104-500ML) and 1% sodium pyruvate (Sartorius, 03-042-1B). Jurkat E6-1 cells were cultured in RPMI 1640 medium (Sartorius, 01-100-1A) supplemented with 10% FBS. Cells were maintained according to the supplier’s recommendations.

### CellTiter-Glo® Luminescent Cell Viability Assay

Cell viability was measured using the CellTiter-Glo luminescent assay (Promega, G7570), according to the manufacturer’s instructions. This assay is based on measuring ATP levels, which indicate metabolically active and viable cells. The luminescence signal is generated by a luciferase reaction and is proportional to the amount of ATP in the sample. Jurkat E6-1 cells were used for this assay. Cells were treated with peptides (TAT–C16 and R11–C16) or cyclosporin A (CsA, Selleckchem, S2286) at different concentrations (50 µM, 20 µM, 10 µM, 1 µM, and 150 nM). Cell viability was measured 48 h after peptide addition. For each condition, approximately 30,000 cells were used per replicate in a total volume of 50 µL, and an equal volume of CellTiter-Glo reagent was added in a 96-well plate. Plates were mixed for 2 min to induce cell lysis and incubated for 10 min at room temperature. Luminescence was measured using an Agilent BioTek Synergy H1 plate reader and recorded as relative light units (RLU). Each condition was measured in triplicate. Cells without treatment were used as control.

### Cell penetration assay

HeLa cells were used to evaluate CPP–peptide penetration. Cells were seeded at 3,000 cells per well in a 96-well glass-bottom plate (Cellvis, P96-1.5H-N) in culture media, 2 days before the experiment. On the day of the experiment, media was removed and replaced with fresh media containing either 0.1% DMSO (cells only control) or TAMRA-labeled peptides (C16, TAT–C16, or R11–C16) at final concentrations of 10 µM or 1 µM. Cells were incubated for 3 h at 37°C in a humidified incubator with 5% CO₂. After incubation, cells were washed to remove excess peptide and fixed with 4% paraformaldehyde (PFA). Fixed cells were stained with DAPI (Abcam, ab228549) to label nuclei. Images were acquired using a ZEISS LSM 900 confocal microscope at 20× magnification. Image analysis was performed using FIJI (ImageJ). Individual cells were manually selected and regions of interest (ROIs) were defined. The mean fluorescence intensity of the TAMRA signal was measured for each cell. For each condition, 30 cells were analyzed. For flow cytometry-based analysis of peptide penetration, HeLa cells were incubated with TAMRA-labeled peptides (C16, TAT–C16, or R11–C16) at final concentrations of 10 µM, 1 µM, or 150 nM for 3 h at 37°C and 5% CO₂. After incubation, cells were washed with PBS and detached using trypsin to remove membrane-bound peptides before analysis. Cells were washed again and analyzed using an S1000EXi flow cytometer. Data were analyzed using FlowJo v11 software. Three independent biological repeats were performed for each condition.

### NFAT nuclear translocation assay

The pcDNA3.1-NFAT1-eGFP plasmid (GenScript) was used for NFAT1 expression studies. Plasmid DNA was prepared using the PureLink™ HiPure Plasmid Midiprep Kit (Thermo Scientific) to obtain high DNA concentration for transfection. HeLa cells were seeded at 3,000 cells per well in 96-well glass-bottom plates 2 days before the experiment. Cells were transfected with the NFAT1-eGFP plasmid using Lipofectamine LTX (Thermo Scientific, 15338030) according to the manufacturer’s instructions. 24 h after transfection, media was replaced with fresh media containing the treatments: DMSO (resting cells and activated cells), 10 µM TAT or R9, 150 nM CsA, TAT–C16 or R11–C16 (at 10 µM, 1 µM, 0.5 µM, 150 nM and 50 nM). Cells were incubated with treatments for 3 h at 37°C and 5% CO₂. Afterwards, cells were stimulated with Phorbol 12-myristate 13-acetate (PMA, Sigma-Aldrich, P8139) and ionomycin (Sigma-Aldrich, I0634) at final concentrations of 10 ng/mL and 1 µM, respectively. Control cells (resting cells) received DMSO. Cells were incubated for an additional 1.5 h. Cells were then washed, fixed with 4% PFA, and stained with DAPI to label nuclei. Images were acquired using a Leica confocal microscope with a 63× oil objective. Image analysis was performed using FIJI (ImageJ). Transfected cells were identified based on GFP signal. Regions of interest (ROIs) were manually defined for the nucleus (based on DAPI staining) and the cytoplasm (based on cell shape in brightfield). Mean GFP fluorescence intensity was measured in both regions. The nuclear-to-cytoplasmic (N/C) fluorescence ratio was calculated for each cell. Data were normalized to the activation control (PMA and ionomycin). For each condition, 30 cells were analyzed per experiment, with three independent biological repeats. Statistical analysis was performed using GraphPad Prism.

### Quantitative real-time PCR

Jurkat E6-1 cells were used to evaluate IL-2 gene expression. For each condition, 1 × 10⁶ cells were seeded per well. Cells were pre-treated for 1.5 h with DMSO (cells only and activation control), 150 nM CsA, or peptides [10 µM TAT or R9, TAT–C16, R11–C16 (10 µM, 1 µM, 0.5 µM, 150 nM and 50 nM)] at 37°C and 5% CO₂. After pre-treatment, cells were stimulated with PMA and ionomycin at final concentrations of 10 ng/mL and 1 µM, respectively, and incubated for an additional 4.5 h. Control cells received DMSO. After a total of 6 h, cells were washed and cell pellets were collected and stored at −80°C until further processing. Total RNA was extracted using the Universal RNA Purification Kit (E3598) according to the manufacturer’s protocol. For each sample, 1 µg RNA was used for cDNA synthesis using the iScript cDNA Synthesis Kit (Bio-Rad, 170-8891). cDNA was diluted to a final volume of 100 µL using ultra-pure water, and 4 µL was used per real-time PCR reaction. Reactions were performed in triplicate using SYBR Green master mix (PB20.16-50) in NEST plates (402001) on a QuantStudio 5 system. RPLP0 was used as a reference gene and IL-2 as the target gene. Primer sequences were obtained from OriGene:

RPLP0 forward: TGGTCATCCAGCAGGTGTTCGA

RPLP0 reverse: ACAGACACTGGCAACATTGCGG

IL-2 forward: AGAACTCAAACCTCTGGAGGAAG

IL-2 reverse: GCTGTCTCATCAGCATATTCACAC

Gene expression was calculated as log₂ fold change and normalized to the activation control (PMA + ionomycin) within each experiment. Each condition included three technical replicates, and the experiment was repeated three times independently. Data were analyzed using GraphPad Prism.

### Luciferase assay

NFAT activity was measured using Jurkat PD-1 luciferase reporter cells (InvivoGen, rajkt-hpd1) and Raji-apc-null cells (InvivoGen, raji-apc-null). Cells were cultured in IMDM medium (Sartorius, 01-058-1A) supplemented with 10% FBS. The assay was performed according to the manufacturer’s protocol with a small modification. Jurkat PD-1 luciferase cells were pre-incubated for 1.5 h with treatments including DMSO (control), 10 µM TAT or R9, TAT–C16 and R11–C16 (10 µM, 1 µM and 150 M) at 37°C and 5% CO₂. After pre-treatment, Raji-apc-null cells were added, and the co-culture was incubated for an additional 4.5 h. The assay was then continued according to the manufacturer’s protocol. Luciferase activity was measured using Bio-Glo reagent (InvivoGen, rep-qlc4lg1) in white 96-well plates (Greiner, 655083). Luminescence was recorded using an Agilent BioTek Synergy H1 plate reader. Each condition was measured in triplicate, and the experiment was repeated three times independently. Data were normalized to the cells only condition and analyzed using GraphPad Prism.

## Results

### Modeling and validation of CPP–C16 binding to Cn

To enable intracellular delivery of the C16 peptide, we conjugated it to TAT and R11 CPPs. To verify that the addition of these cationic sequences does not compromise the peptide’s interaction with Cn, we first performed structural modeling of the TAT–C16 and R11–C16 conjugates (CPP-C16) in complex with the CnA subunit. The resulting models predict that the core C16 peptides bind to CnA, while the CPP extensions remain solvent-exposed without sterically interfering with the binding interface (Fig. 2A). We next experimentally validated the binding of the conjugated peptides using a fluorescence polarization (FP) competition assay. In this assay, the TAT-C16 and R11-C16 peptides were titrated against a pre-formed complex of CnA and fluorescein-labeled PVIVIT (FITC-PVIVIT), a high-affinity ligand used as a probe [15,35]. In the absence of competitors, the large FITC-PVIVIT/CnA complex exhibits high polarization values due to the restricted rotation of the fluorophore. Upon titration, both TAT–C16 and R11–C16 induced a dose-dependent decrease in polarization (Fig. 2B), indicating the successful displacement of the FITC-PVIVIT probe from the CnA interface. The calculated IC50s (Table 1) confirm that the CPP-conjugated peptides retain a potent binding affinity for Cn that is strictly comparable to the non-conjugated C16 peptide.

**Figure 2.**
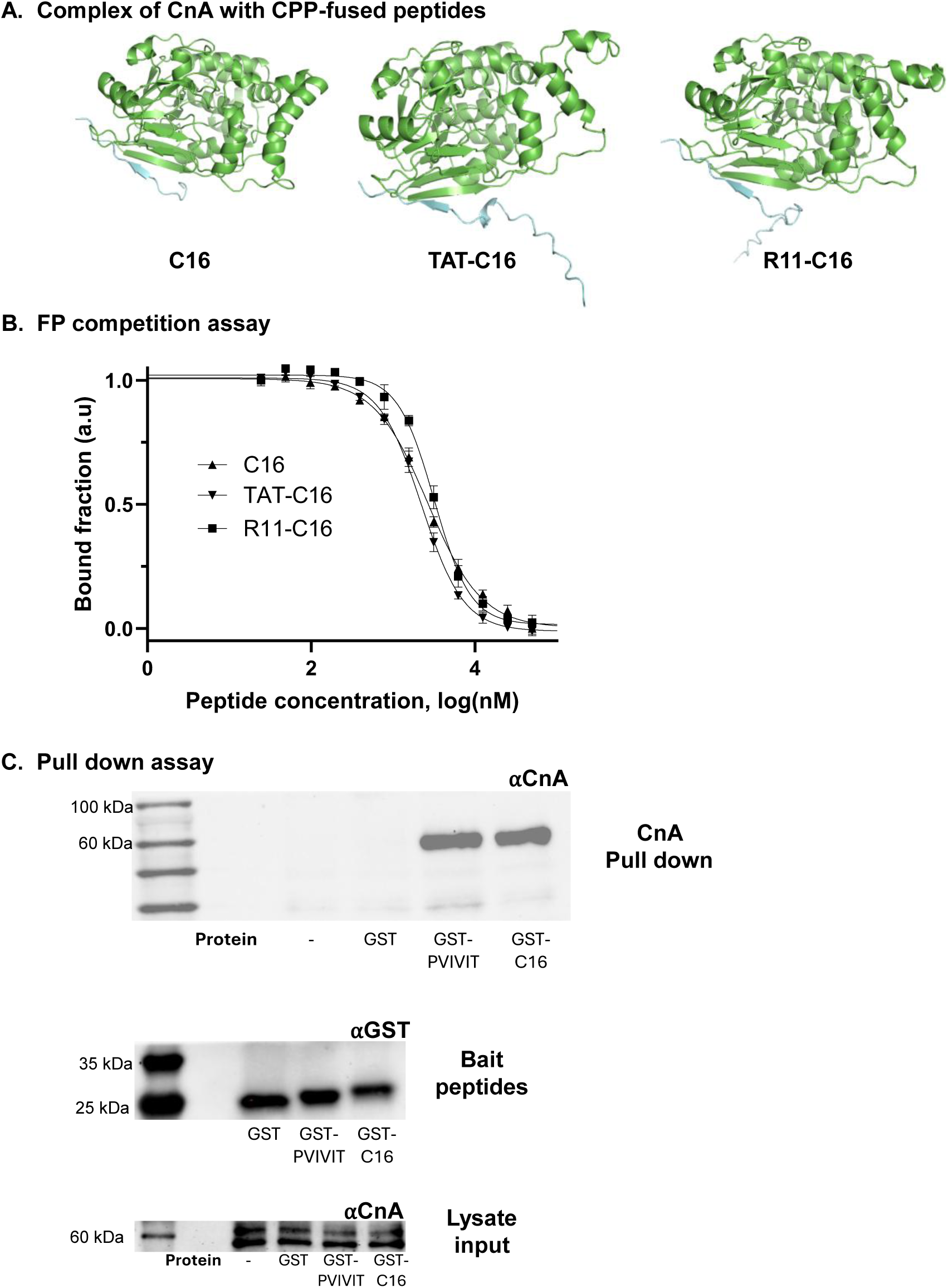
Structural and biophysical characterization of C16 peptide variants binding to CnA. **(A)** Structural models illustrating the predicted binding of C16 (left), TAT-C16 (middle), and R11-C16 (right) peptides (cyan) to the CnA subunit (green). The models demonstrate that the conjugation of the CPPs does not sterically hinder the C16 interaction interface. **(B)** FP competition assay. The binding affinities of the three peptide variants were evaluated by their ability to competitively displace a fluorescent probe (FITC-PVIVIT) from CnA. All variants exhibited comparable IC50 values (summarized in Table 1). Data are presented as mean ± SD (n = 3). **(C)** GST pull-down assay validating direct interaction between the C16 peptide and endogenous CnA. (Top panel) Immunoblot showing binding of CnA (∼60 kDa) by GST-C16 and the positive control GST-PVIVIT. (Middle panel) Bait loading control confirming immobilization of the recombinant GST fusion proteins. (Bottom panel) Input control of endogenous CnA across the crude Jurkat cell lysate samples prior to incubation with the beads. Legend: (–) - beads alone; GST – GST protein only; GST-C16 – GST protein fused with C-terminus C16 peptide; GST-PVIVIT – GST protein fused with C-terminus PVIVIT peptide.

**Table 1.** Competitive inhibitory concentrations (IC50) of CPP–C16 peptides against the CnA/FITC-PVIVIT complex.

| Peptide name | IC<sub>50</sub> (μM) |
| --- | --- |
| C16 | 1.37 ± 0.19 |
| R11-C16 | 1.88 ± 0.20 |
| TAT-C16 | 1.18 ± 0.13 |

To evaluate peptide binding to CnA within the complex physiological environment, we verified the ability of the C16 peptide to bind endogenous Cn from cellular lysates. Recombinant GST-tagged C16 peptide was immobilized onto glutathione Sepharose beads, and incubated with native Jurkat cell lysates. Western blot analysis of the pull-down fractions revealed that GST-C16 robustly captured endogenous CnA (∼60 kDa), demonstrating a functional binding efficiency comparable to the canonical high-affinity positive control, GST-PVIVIT (Fig. 2C). No non-specific CnA binding was detected in the negative control groups, which included beads only (–) and GST-alone lanes. Successful bait immobilization and equal cellular protein input across the experimental groups were confirmed via anti-GST and anti-CnA input immunoblots, respectively.

### C16-conjugated to CPP enters mammalian cells with low toxicity

Having confirmed that the CPP–C16 variants retain high affinity for CnA *in vitro* and successfully bind native CnA, we next assessed their cellular permeability. Confocal microscopy of HeLa cells incubated with N-terminus TAMRA-labeled TAT-C16 or R11-C16 revealed entry of the peptides into the cells (Fig. 3A, Fig. S1). Following three hours of treatment, the cells exhibited robust intracellular fluorescence, indicating successful transduction of the peptides across the plasma membrane. In contrast, untreated control cells and TAMRA-labeled C16 peptide without a CPP fusion showed no signal, confirming that the observed fluorescence was driven by peptide uptake rather than nonspecific background. Quantification of the fluorescence signal further validated these observations, with both conjugates demonstrating significant cellular entry (Fig. 3B). To further quantify peptide uptake, cells were analyzed by flow cytometry following incubation with the indicated TAMRA-labeled peptides and peptide internalization was evaluated by mean fluorescence intensity (MFI). The analysis demonstrated the cellular uptake of both CPP-conjugated peptides, whereas untreated cells and cells incubated with TAMRA-C16 alone showed minimal fluorescence signal (Fig. 3C-D). We next evaluated the cytotoxicity of the peptide variants relative to the clinical standard CsA. Jurkat E6-1 cells were treated with increasing concentrations of the peptides (up to 50 µM), and cell viability was subsequently quantified (Fig. 3E). While CsA imparted high toxicity, causing near-complete cell death at 10 µM, the R11-C16 peptide and TAT-C16 exhibited minimal cytotoxicity across the tested dose range. Notably, cell viability remained higher than 75% even at 50 µM.

**Figure 3.**
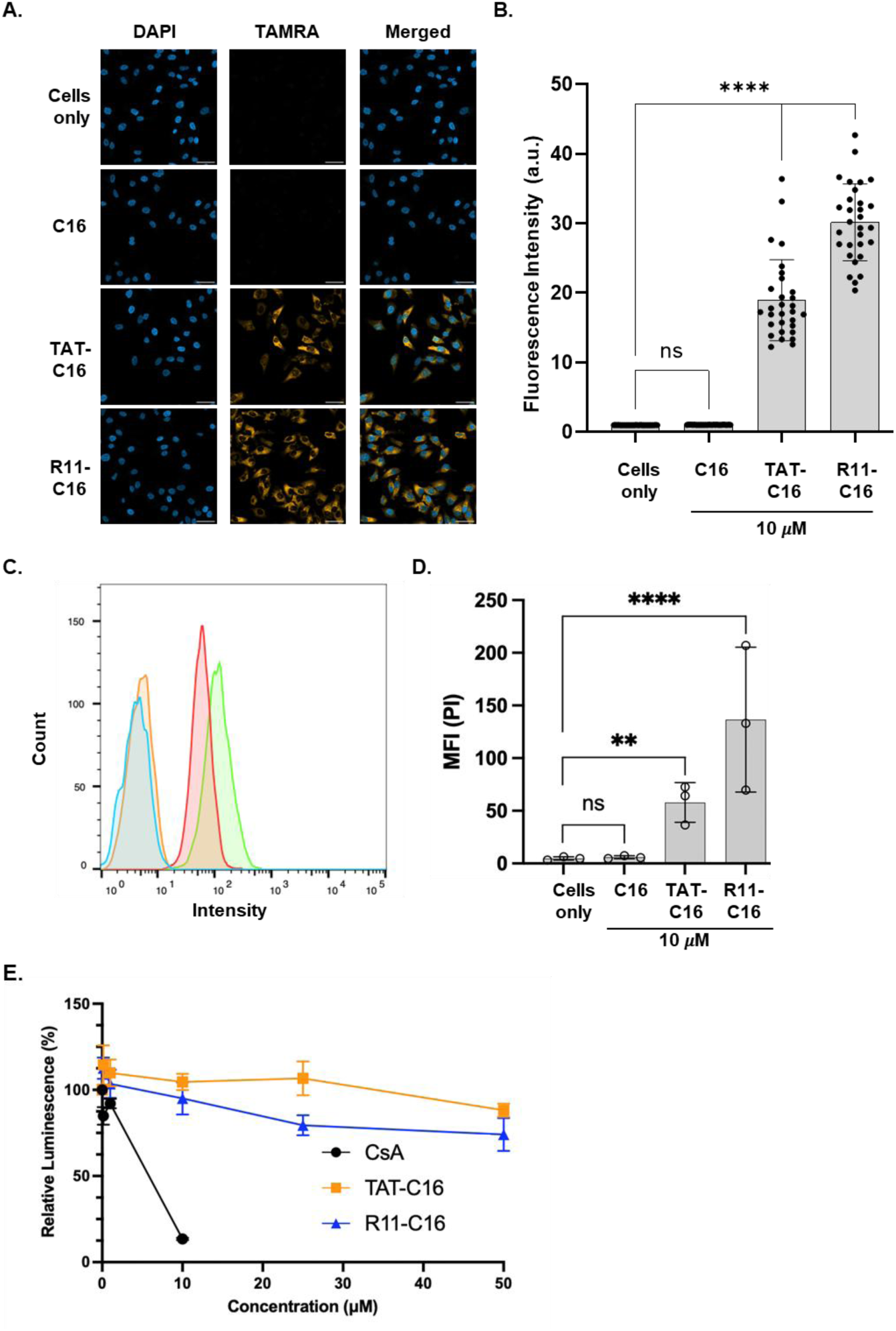
Cellular uptake and cytotoxicity profile of CPP–C16 conjugates. **(A)** HeLa cells were treated with 10 µM of TAMRA-labeled C16, TAT-C16 or R11-C16 peptides. Confocal fluorescence microscopy reveals robust intracellular accumulation of both CPP-conjugated variants compared to untreated control cells ("Cells only") or cells treated with non-labeled C16. Scale bar, 50 µm. **(B)** Quantification of intracellular fluorescence intensity. Individual cells were manually selected, and the mean TAMRA fluorescence intensity was measured for each cell. For each condition, 30 cells were analyzed. The bar plot shows significantly higher cellular uptake for R11–C16 compared to TAT–C16. Statistical significance was determined using one-way ANOVA followed by Dunnett’s multiple comparisons test ****P<0.0001; ns – not significant. **(C)** Representative flow cytometry histograms showing TAMRA fluorescence intensity in HeLa cells following 3 h incubation with TAMRA-labeled CPP–peptides at 10 µM. Blue- cells only, orange- C16, red- TAT-c16 and green- R11-C16. **(D)** Quantification of flow cytometry mean fluorescence intensity (MFI) from peptide-treated HeLa cells. Cells were incubated with TAMRA-labeled peptides for 3 h, washed, trypsinized, and analyzed by flow cytometry. Data are presented as mean ± SEM from three independent biological experiments. Statistical significance was determined using one-way ANOVA followed by Dunnett’s multiple comparisons test **p < 0.01, ****p < 0.0001, ns - not significant. **(E)** Cellular viability of Jurkat T cells following a 48-hour treatment with increasing concentrations (0.1 – 50 µM) of CsA, R11-C16 and TAT-C16. Cell viability was quantified relative to untreated controls. Data are presented as mean ± SD (n = 3).

### CPP-C16 Conjugates Inhibit NFAT Nuclear Translocation

To evaluate the intracellular activity of the CPP-C16 conjugates, we monitored the subcellular localization of GFP-tagged NFAT1 in HeLa cells treated with variable peptide concentrations using fluorescence microscopy (Fig. 4; Fig. S2). In resting cells, GFP-NFAT1 was primarily located in the cytoplasm. Following cellular activation with ionomycin and PMA, we observed robust nuclear translocation of GFP-NFAT1. As a positive inhibitory control, CsA treatment effectively inhibited NFAT nuclear translocation, maintaining NFAT predominantly in the cytosol of activated cells. Treatment with 10 µM of either the TAT-C16 or R11-C16 significantly impaired NFAT nuclear entry. Quantitative analysis of the imaging data revealed that both CPP-C16 conjugates reduced the proportion of cells exhibiting nuclear NFAT to approximately 55–60% compared with untreated activated cells. The data confirm that the CPP-mediated intracellular delivery of the C16 peptide inhibits the Cn-NFAT interaction and retains NFAT in the cytoplasm.

**Figure 4.**
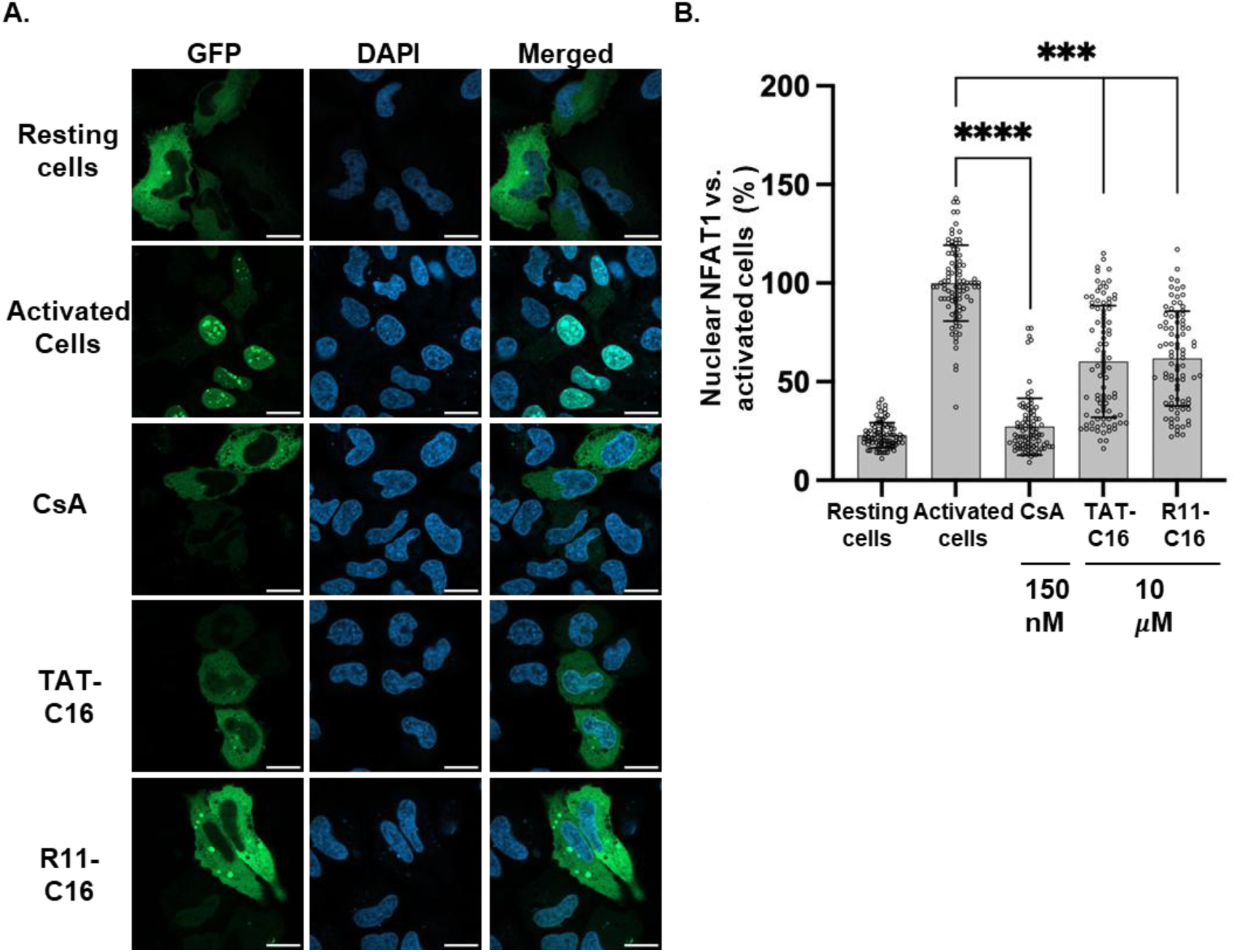
R11-C16 and TAT-C16 conjugates inhibit NFAT1 nuclear translocation. **(A)** Representative confocal fluorescence images of HeLa cells expressing GFP-tagged NFAT1. Stimulation with PMA and ionomycin (activated cells) induces robust nuclear translocation of GFP-NFAT1 (top row). Treatment with the positive control CsA (150 nM), or the peptide conjugates R11-C16 or TAT-C16 (10 µM) effectively inhibits this nuclear entry, retaining GFP-NFAT1 within the cytoplasm. Scale bar = 20 µm. **(B)** Quantification of NFAT1 nuclear entry, expressed as the percentage of cells exhibiting nuclear localization relative to the untreated, activated control cells (100%). Data are presented as mean ± SD of independent biological replicates ****P<0.0001, ***P<0.001.

### CPP-C16 conjugates inhibit NFAT-driven transcription and *IL-2* expression in human T cells

Having established that the CPP-C16 conjugates successfully inhibit the nuclear translocation of NFAT, we next sought to determine whether this cytosolic retention translates to a reduction in downstream NFAT-dependent transcriptional activity. To this end, we utilized a functional immune co-culture bioassay comprising Raji antigen-presenting cells and Jurkat T cells stably expressing a luciferase reporter gene under the control of an NFAT-response element. Upon cellular activation of Jurkat cells via T-cell receptor (TCR) engagement, we observed a robust induction of NFAT-driven luciferase luminescence (Fig. 5A). As expected, treatment with the clinical standard CsA reduced the signal to 30% of that observed in activated, vehicle-treated cells. Similarly, treatment with 10 µM of either R11-C16 or TAT-C16 resulted in a significant reduction in luciferase activity, reducing the signal to approximately 70% of the activated control. Treatment with only CPPs did not affect luciferase activity (Fig. S3A).

**Figure 5.**
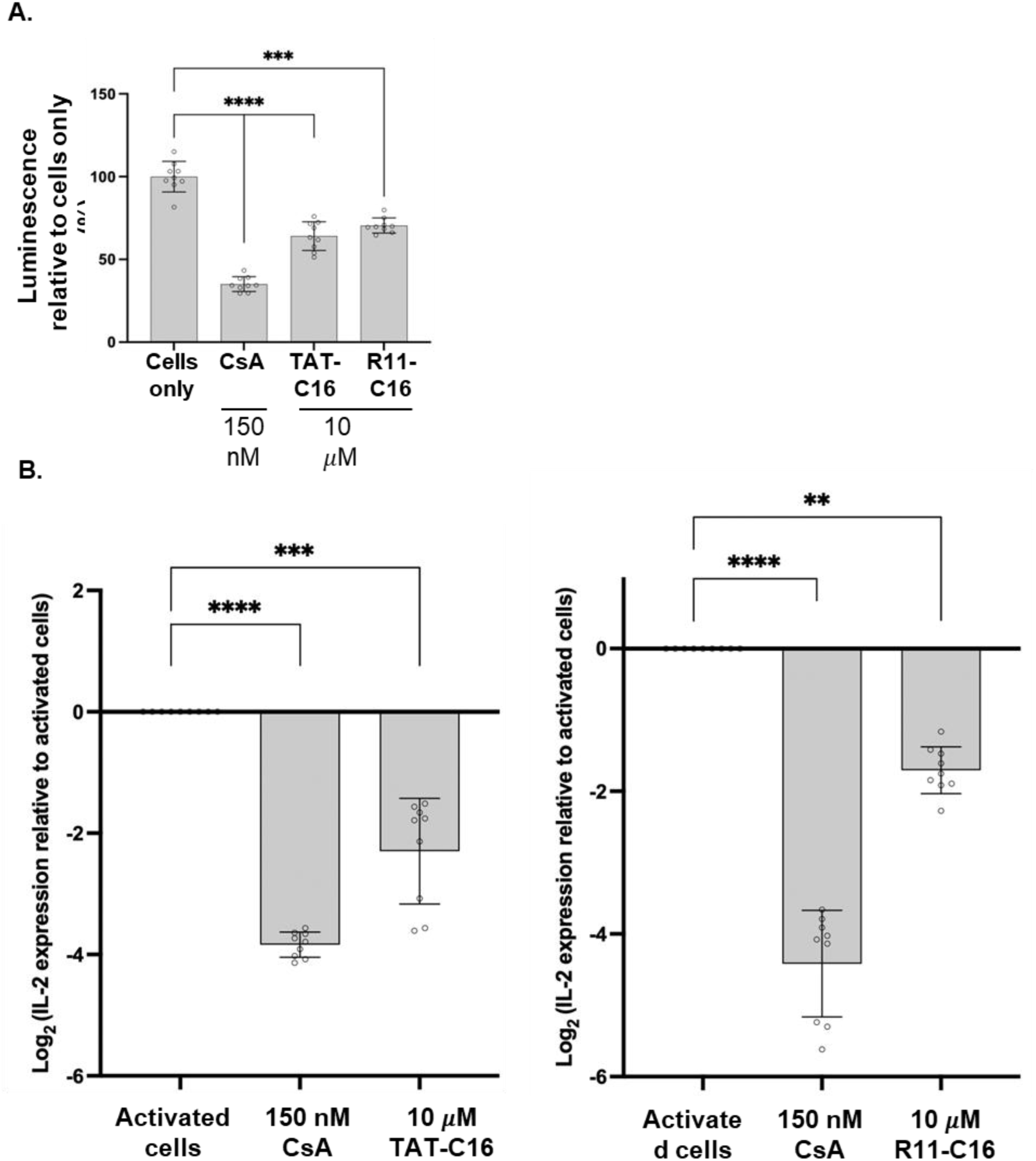
R11-C16 and TAT-C16 peptides inhibit NFAT-dependent transcriptional activity and *IL-2* levels in human T cells. **(A)** Evaluation of NFAT transcriptional activity using an immune co-culture bioassay. Jurkat T cells stably expressing a luciferase reporter under the control of an NFAT-response element were co-cultured with Raji antigen-presenting cells. Cells were treated with vehicle (activated cells), the positive control CsA (150 nM), or the peptide conjugates R11-C16 or TAT-C16 (10 µM). Luminescence was measured and is expressed relative to the activated control. Data are presented as mean ± SD of independent replicates. ****P<0.001; ***P<0.005; ns – non significant. **(B)** Quantitative RT-PCR analysis of IL-2 gene expression in Jurkat T cells. Cells were pre-treated with R11-C16 or CsA for 1.5 hours, followed by stimulation with PMA and ionomycin for 6 hours. Results are expressed as log fold change relative to unstimulated cells ("Cells only"). Data are presented as mean ± SD. ****P<0.001; ***P<0.005; ns – non significant.

To further validate these findings in the context of endogenous immune signaling, we measured the transcription of Interleukin-2 (IL-2), a primary downstream target of the Cn-NFAT pathway, using quantitative real-time PCR (qRT-PCR) (Fig. 5B). Stimulation of Jurkat cells induced an upregulation of IL-2 mRNA levels compared to the resting state. Treatment with 150 nM CsA profoundly suppressed this expression. In accordance with the reporter assay results, treatment with 10 µM of R11-C16 or TAT-C16 significantly attenuated the transcriptional upregulation of IL-2 in the activated cells. Treatment with only CPPs did not affect IL-2 levels (Fig. S3B). These data confirm that CPP-C16-mediated cytosolic retention of NFAT effectively impairs its transcriptional activity, ultimately preventing the expression of key T-cell-activating cytokines.

## Discussion

Inhibition of PPIs represents a significant opportunity in modern drug discovery, offering a distinct advantage over traditional catalytic-site blockade. While inhibitors such as the clinical standard CsA effectively block enzymatic activity, affecting multiple molecular pathways, which in turn translates to clinical toxicity. In contrast, targeting the Cn-NFAT PPI interface, such as the specific PxIxIT docking surface in Cn, offers a potential strategy to decouple enzyme-catalytic activity from PPI inhibition, potentially yielding efficacy without toxicity. Whereas efforts have been made to develop small-molecule Cn-NFAT PPI inhibitors [33], the challenging Cn-NFAT interface rendered these less effective than peptides. Central to the study of selective Cn–NFAT peptide inhibitors is the synthetic PVIVIT peptide, an engineered, high-affinity variant optimized from the native consensus PxIxIT motif in NFAT [24,26]. Various studies have successfully incorporated the PVIVIT peptide, establishing it as the benchmark tool for substrate-specific Cn blockade. Cellular activity of the PVIVIT peptide was achieved by conjugating it to a polyarginine delivery vector (11R-VIVIT) [28]. Subsequent structure-based optimization has yielded PVIVIT variants and peptidomimetics with dissociation constants in the nanomolar range, significantly enhancing their ability to compete with the binding of native NFAT with Cn and with improved pharmacological properties [25,34]. Furthermore, connecting PxIxIT- and LxVP-targeting moieties via a flexible linker have proven highly effective in modulating the Cn-NFAT signaling and the addition of a CPP conjugation established their bioactivity in cells and in vivo [35–37]. The ability for intracellular inhibition of the Cn-NFAT interaction has led to various biomedical applications, some beyond the classic T-cells system [27,34,38–40].

Herein, we evaluated the ability to target the Cn-NFAT PPI and have shown that the TAT-C16 and R11–C16 fusion peptides act as effective inhibitors of this interaction. Structural modeling and biochemical assays confirmed that the fusion of cell-penetrating motifs does not compromise the peptide’s affinity for the PxIxIT docking epitope on Cn. Functionally, these peptides successfully translocate across the plasma membrane to directly reduce NFAT nuclear translocation and subsequent IL-2 gene expression, mimicking the immunosuppressive phenotype of CsA. Importantly, this efficacy was achieved through enhanced cellular viability conferred by the peptide. While CsA exhibited significant cytotoxicity at concentrations as low as 1 μM, the peptides remained non-toxic even at 50 μM, establishing a favorable therapeutic window that is often elusive for small-molecule inhibitors in general and CsA in specific. Our study therefore demonstrates that a substrate-competitive peptide can achieve a desired biological effect based on PPI inhibition without the systemic toxicity associated with catalytic blockade.

Despite these promising in vitro results, translating linear peptides such as CPP–C16 into clinical therapeutics poses inherent challenges. Linear peptides are susceptible to rapid degradation by serum proteases and intracellular peptidases, resulting in a short half-life in vivo. Furthermore, the flexibility of the linear backbone imposes an entropic penalty upon binding, which can limit the maximal affinity and selectivity required for a drug candidate. Consequently, while CPP–C16 serves as an excellent chemical probe for validating the mechanism, its utility as a systemic drug may be limited by poor pharmacokinetics and the need for high dosing frequencies to maintain therapeutic levels. To overcome these barriers, future research must focus on structural optimization to enhance metabolic stability and bioavailability. A primary strategy is peptide cyclization or stapling. This could constrain the peptide into the bioactive conformation and, most importantly, protect it from proteolytic cleavage. Such a molecule would likely retain the high affinity and safety profile of PPI modulator but with the pharmacokinetic durability required for in-vivo application.

## Conclusion

We demonstrated that conjugating the novel C16 peptide with cell-penetrating motifs effectively overcomes the membrane barrier, enabling robust intracellular delivery and specific inhibition of NFAT nuclear translocation. This targeted blockade successfully dampens the immune response without severe cytotoxicity. While the linear CPP–C16 peptide serves as a powerful proof-of-concept, translating this strategy into a clinically viable immunosuppressant will require the development of stable, cyclized peptidomimetics.

## Materials and Methods

### Peptides

All peptides were synthesized with C-terminal amidation by GL Biochem (Shanghai, China). Peptides were dissolved in DMSO and diluted to working concentrations before use. Peptide sequences were the following: (C16) KHLDVPDIIITPPTPT; (TAT-C16) GRKKRRQRRRPQKHLDVPDIIITPPTPT and (R11-C16) RRRRRRRRRRRKHLDVPDIIITPPTPT.

### GST-fusion peptides expression and purification

GST, GST-PVIVIT and GST-C16 fusion proteins were expressed in *Escherichia coli* BL21 cells as described earlier [15]. Briefly, bacterial cultures were grown to an OD600 of approximately 0.8 and protein expression was induced with 1 mM IPTG overnight at 25°C. Bacterial pellets were lysed in PBS containing protease inhibitor (Merck, 11836170001), and GST-tagged proteins were purified using an AKTA purification system with a GSTrap HP column (Cytiva). Purified proteins were dialyzed against PBS, centrifuged at 18,000 × g for 20 min to remove aggregates, and protein concentration was determined using NanoDrop spectrophotometry and BCA assay (Thermo fisher, 23225).

### GST pull-down assay

For pull-down experiments, 8 × 10^7^ Jurkat E6-1 cells were collected, washed with PBS, and lysed in RIPA buffer containing 1:100 protease inhibitor. For each condition, 30 µL Glutathione Sepharose 4B beads (Cytiva, 17-0756-01) were washed and incubated with 25 µg GST-tagged proteins or beads only as a negative control for 2 h at 4°C on a rotator. After removal of unbound proteins, Jurkat cell lysates were added to the beads and incubated for 3 h at 4°C on a rotator. Beads were then washed with RIPA buffer containing protease inhibitor, and bound proteins were eluted using 4× sample buffer followed by boiling at 95°C for 10 min prior to western blot analysis.

### Western blotting

Proteins were separated by SDS-PAGE using 12% acrylamide gels (1.5 mm thickness) and transferred onto PVDF membranes (Bio-Rad, 1704157). Membranes were then probed for detection of GST-tagged proteins using a mouse anti-GST antibody (GenScript, A00865) and for CnA using a rabbit anti-CnA antibody (Abcam, ab52761). HRP-conjugated secondary antibodies were used for detection: goat anti-mouse HRP (Sigma-Aldrich, AP130P) for mouse primary antibody and goat anti-rabbit HRP (Sigma-Aldrich, 12-348) for rabbit primary antibody.

### Cells and culture conditions

HeLa cells (ATCC, CCL-2) and Jurkat E6-1 cells (ATCC, TIB-152) were used in this study. Cells were maintained at 37°C in a humidified incubator with 5% CO₂. HeLa cells were cultured in high-glucose DMEM (Sartorius, 01-052-1A) supplemented with 10% fetal bovine serum (FBS, Diagnovum, D104-500ML) and 1% sodium pyruvate (Sartorius, 03-042-1B). Jurkat E6-1 cells were cultured in RPMI 1640 medium (Sartorius, 01-100-1A) supplemented with 10% FBS. Cells were maintained according to the supplier’s recommendations.

### CellTiter-Glo® Luminescent Cell Viability Assay

Cell viability was measured using the CellTiter-Glo luminescent assay (Promega, G7570), according to the manufacturer’s instructions. This assay is based on measuring ATP levels, which indicate metabolically active and viable cells. The luminescence signal is generated by a luciferase reaction and is proportional to the amount of ATP in the sample. Jurkat E6-1 cells were used for this assay. Cells were treated with peptides (TAT–C16 and R11–C16) or cyclosporin A (CsA, Selleckchem, S2286) at different concentrations (50 µM, 20 µM, 10 µM, 1 µM, and 150 nM). Cell viability was measured 48 h after peptide addition. For each condition, approximately 30,000 cells were used per replicate in a total volume of 50 µL, and an equal volume of CellTiter-Glo reagent was added in a 96-well plate. Plates were mixed for 2 min to induce cell lysis and incubated for 10 min at room temperature. Luminescence was measured using an Agilent BioTek Synergy H1 plate reader and recorded as relative light units (RLU). Each condition was measured in triplicate. Cells without treatment were used as control.

### Cell penetration assay

HeLa cells were used to evaluate CPP–peptide penetration. Cells were seeded at 3,000 cells per well in a 96-well glass-bottom plate (Cellvis, P96-1.5H-N) in culture media, 2 days before the experiment. On the day of the experiment, media was removed and replaced with fresh media containing either 0.1% DMSO (cells only control) or TAMRA-labeled peptides (C16, TAT–C16, or R11–C16) at final concentrations of 10 µM or 1 µM. Cells were incubated for 3 h at 37°C in a humidified incubator with 5% CO₂. After incubation, cells were washed to remove excess peptide and fixed with 4% paraformaldehyde (PFA). Fixed cells were stained with DAPI (Abcam, ab228549) to label nuclei. Images were acquired using a ZEISS LSM 900 confocal microscope at 20× magnification. Image analysis was performed using FIJI (ImageJ). Individual cells were manually selected and regions of interest (ROIs) were defined. The mean fluorescence intensity of the TAMRA signal was measured for each cell. For each condition, 30 cells were analyzed. For flow cytometry-based analysis of peptide penetration, HeLa cells were incubated with TAMRA-labeled peptides (C16, TAT–C16, or R11–C16) at final concentrations of 10 µM, 1 µM, or 150 nM for 3 h at 37°C and 5% CO₂. After incubation, cells were washed with PBS and detached using trypsin to remove membrane-bound peptides before analysis. Cells were washed again and analyzed using an S1000EXi flow cytometer. Data were analyzed using FlowJo v11 software. Three independent biological repeats were performed for each condition.

### NFAT nuclear translocation assay

The pcDNA3.1-NFAT1-eGFP plasmid (GenScript) was used for NFAT1 expression studies. Plasmid DNA was prepared using the PureLink™ HiPure Plasmid Midiprep Kit (Thermo Scientific) to obtain high DNA concentration for transfection. HeLa cells were seeded at 3,000 cells per well in 96-well glass-bottom plates 2 days before the experiment. Cells were transfected with the NFAT1-eGFP plasmid using Lipofectamine LTX (Thermo Scientific, 15338030) according to the manufacturer’s instructions. 24 h after transfection, media was replaced with fresh media containing the treatments: DMSO (resting cells and activated cells), 10 µM TAT or R9, 150 nM CsA, TAT–C16 or R11–C16 (at 10 µM, 1 µM, 0.5 µM, 150 nM and 50 nM). Cells were incubated with treatments for 3 h at 37°C and 5% CO₂. Afterwards, cells were stimulated with Phorbol 12-myristate 13-acetate (PMA, Sigma-Aldrich, P8139) and ionomycin (Sigma-Aldrich, I0634) at final concentrations of 10 ng/mL and 1 µM, respectively. Control cells (resting cells) received DMSO. Cells were incubated for an additional 1.5 h. Cells were then washed, fixed with 4% PFA, and stained with DAPI to label nuclei. Images were acquired using a Leica confocal microscope with a 63× oil objective. Image analysis was performed using FIJI (ImageJ). Transfected cells were identified based on GFP signal. Regions of interest (ROIs) were manually defined for the nucleus (based on DAPI staining) and the cytoplasm (based on cell shape in brightfield). Mean GFP fluorescence intensity was measured in both regions. The nuclear-to-cytoplasmic (N/C) fluorescence ratio was calculated for each cell. Data were normalized to the activation control (PMA and ionomycin). For each condition, 30 cells were analyzed per experiment, with three independent biological repeats. Statistical analysis was performed using GraphPad Prism.

### Quantitative real-time PCR

Jurkat E6-1 cells were used to evaluate IL-2 gene expression. For each condition, 1 × 10⁶ cells were seeded per well. Cells were pre-treated for 1.5 h with DMSO (cells only and activation control), 150 nM CsA, or peptides [10 µM TAT or R9, TAT–C16, R11–C16 (10 µM, 1 µM, 0.5 µM, 150 nM and 50 nM)] at 37°C and 5% CO₂. After pre-treatment, cells were stimulated with PMA and ionomycin at final concentrations of 10 ng/mL and 1 µM, respectively, and incubated for an additional 4.5 h. Control cells received DMSO. After a total of 6 h, cells were washed and cell pellets were collected and stored at −80°C until further processing. Total RNA was extracted using the Universal RNA Purification Kit (E3598) according to the manufacturer’s protocol. For each sample, 1 µg RNA was used for cDNA synthesis using the iScript cDNA Synthesis Kit (Bio-Rad, 170-8891). cDNA was diluted to a final volume of 100 µL using ultra-pure water, and 4 µL was used per real-time PCR reaction. Reactions were performed in triplicate using SYBR Green master mix (PB20.16-50) in NEST plates (402001) on a QuantStudio 5 system. RPLP0 was used as a reference gene and IL-2 as the target gene. Primer sequences were obtained from OriGene:

RPLP0 forward: TGGTCATCCAGCAGGTGTTCGA

RPLP0 reverse: ACAGACACTGGCAACATTGCGG

IL-2 forward: AGAACTCAAACCTCTGGAGGAAG

IL-2 reverse: GCTGTCTCATCAGCATATTCACAC

Gene expression was calculated as log₂ fold change and normalized to the activation control (PMA + ionomycin) within each experiment. Each condition included three technical replicates, and the experiment was repeated three times independently. Data were analyzed using GraphPad Prism.

### Luciferase assay

NFAT activity was measured using Jurkat PD-1 luciferase reporter cells (InvivoGen, rajkt-hpd1) and Raji-apc-null cells (InvivoGen, raji-apc-null). Cells were cultured in IMDM medium (Sartorius, 01-058-1A) supplemented with 10% FBS. The assay was performed according to the manufacturer’s protocol with a small modification. Jurkat PD-1 luciferase cells were pre-incubated for 1.5 h with treatments including DMSO (control), 10 µM TAT or R9, TAT–C16 and R11–C16 (10 µM, 1 µM and 150 M) at 37°C and 5% CO₂. After pre-treatment, Raji-apc-null cells were added, and the co-culture was incubated for an additional 4.5 h. The assay was then continued according to the manufacturer’s protocol. Luciferase activity was measured using Bio-Glo reagent (InvivoGen, rep-qlc4lg1) in white 96-well plates (Greiner, 655083). Luminescence was recorded using an Agilent BioTek Synergy H1 plate reader. Each condition was measured in triplicate, and the experiment was repeated three times independently. Data were normalized to the cells only condition and analyzed using GraphPad Prism.

## CRediT authorship contributions

**Adi Cohen:** Methodology, Investigation, Data curation, Writing – original draft, Writing – review & editing; **Matan Gabay:** Methodology, Investigation, Data curation. **Sumit Gupta:** Methodology, Investigation; **Marina Sova:** Methodology, Investigation, Project administration. **Daniel Bar:** Methodology, Writing – review & editing. **Jerome Tubiana:** Conceptualization, Supervision, Writing – original draft, Writing – review & editing. **Maayan Gal:** Conceptualization, Supervision, Funding acquisition, Writing – original draft, Writing – review & editing.

## Data availability

The data is available upon request from the authors.

## Acknowledgment

This research was supported by funding from Colton Center for Autoimmune Diseases (Tel Aviv University), Israeli Ministry of Innovation, Science and Technology, Israeli Innovation Authority (IIA) #89393. Adi Cohen is grateful for the Ph.D. scholarship financial support from the Longevity and Health Research Center at Tel Aviv University. Matan Gabay is grateful to the Marian Gertner Institute for Medical Nano systems and the Yoran Institute for Advanced PhD Students in Personalized Medicine.

## Ethical statement - studies in humans and animals

This study does not involve human or animal subjects.

