## Supplementary material for "A fusion Cell-Permeable C16orf74 Peptide Selectively Disrupts Calcineurin-NFAT Interaction and Inhibits T-cell Activation Without Cytotoxicity": Fig. S

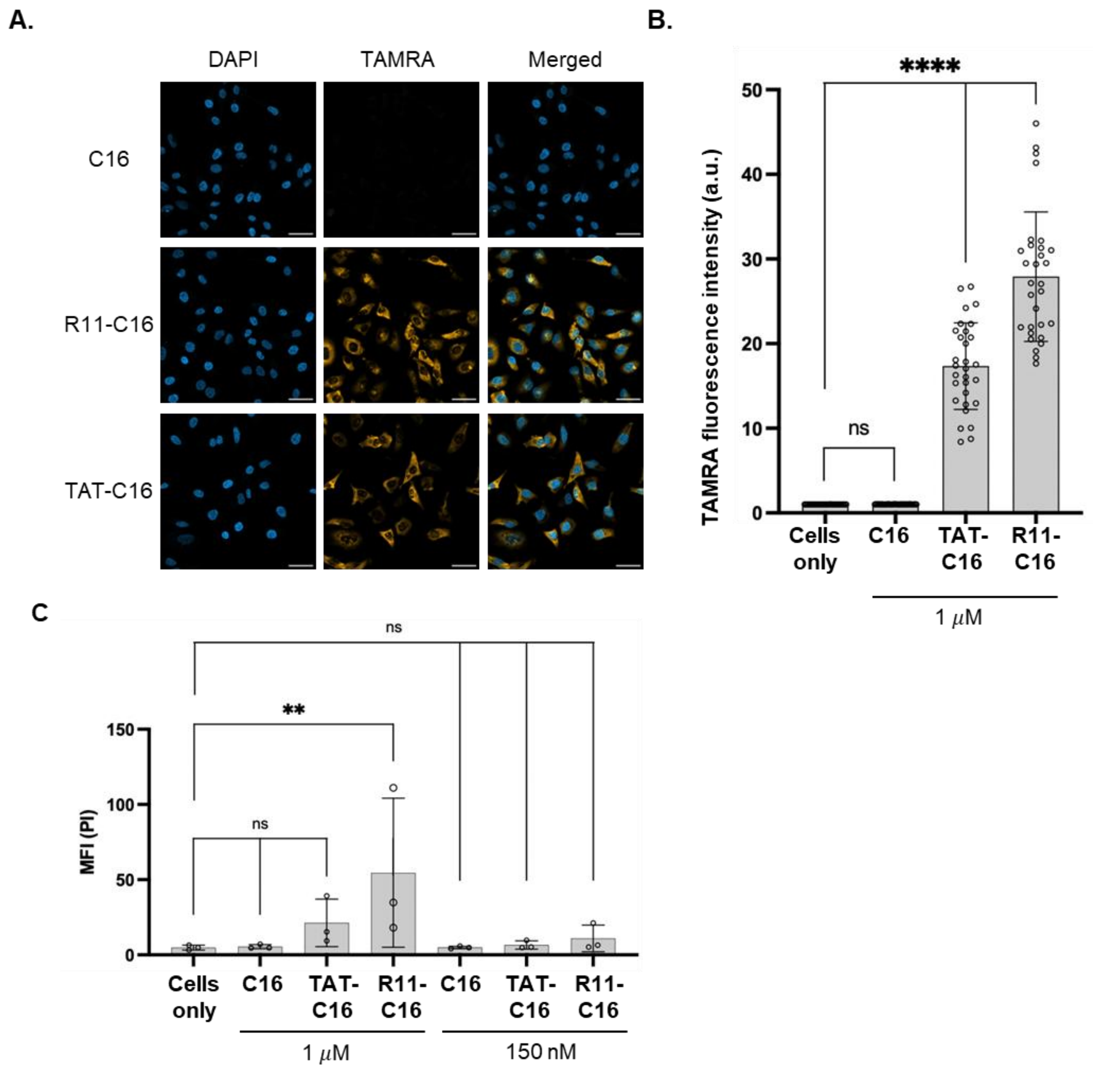


**Figure S1. Cellular permeability of TAT- and R11-C16. (A)** HeLa cells were treated with the 1 µM of TAMRA-labeled C16, TAT-C16 or R11-C16 peptides. Confocal fluorescence microscopy reveals robust intracellular accumulation of both CPP-conjugated variants compared to untreated control cells ("Cells only") or cells treated with C16. Scale bar, 50 µm. (B) Quantification of intracellular fluorescence intensity. Individual cells were manually selected, and the mean TAMRA fluorescence intensity was measured for each cell. For each condition, 30 cells were analyzed. The bar plot shows significantly higher cellular uptake for R11–C16 compared to TAT–C16. ****P<0.0001; ns – not significant. (C) Quantification of intracellular fluorescence intensity of cells treated with variable concentrations of the CPPs alone and the CPP-C16 peptide. **P<0.01; ns – not significant

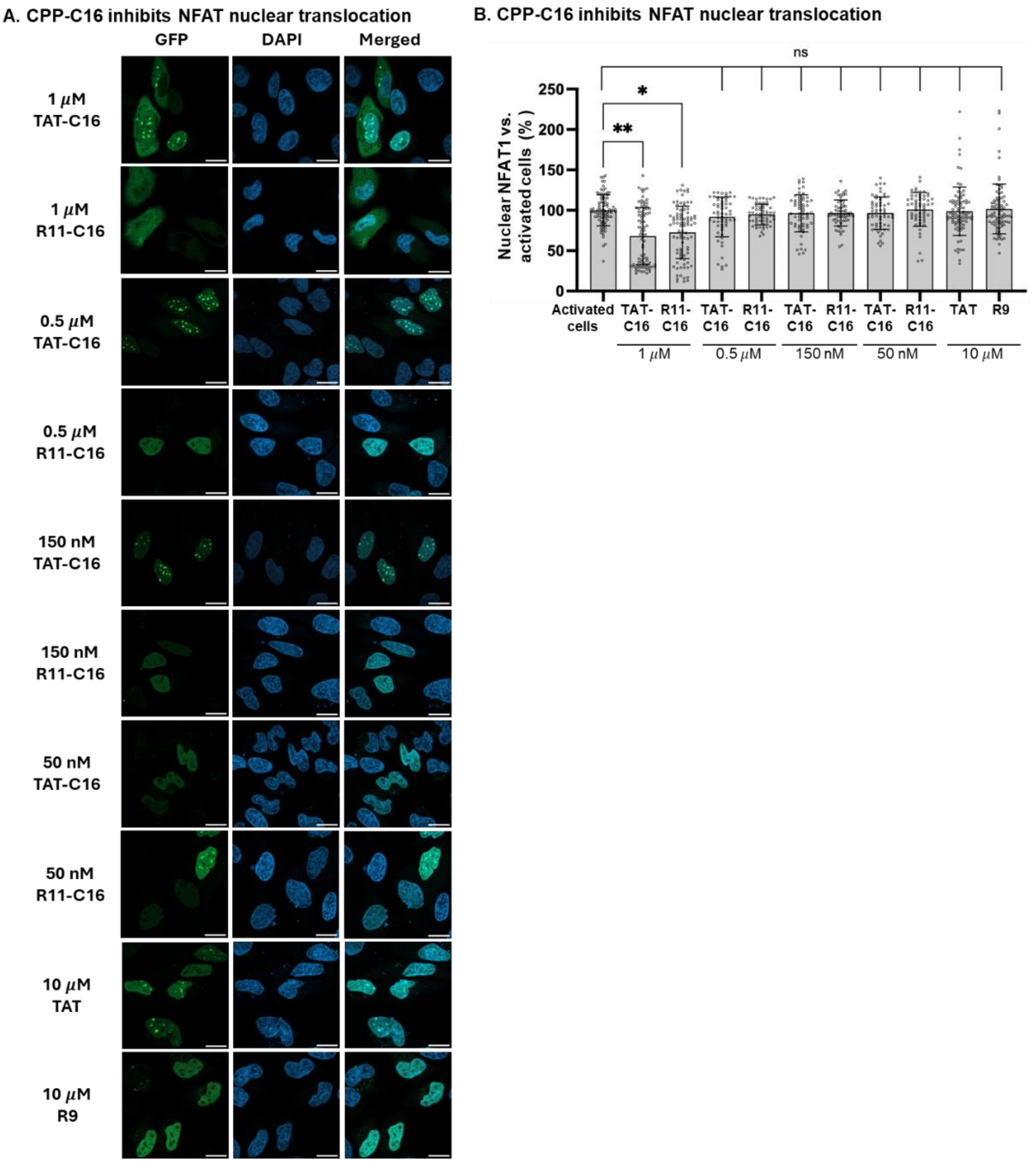


**Figure S2. R11-C16 and TAT-C16 conjugates inhibit NFAT1 nuclear translocation. (A)** Representative confocal fluorescence images of HeLa cells expressing GFP-tagged NFAT1. Stimulation with PMA and ionomycin (activated cells) induces robust nuclear translocation of GFP-NFAT1 (top row). Treatment with the positive control CsA (150 nM), or the peptide conjugates R11-C16 or TAT-C16 at variable concentrations showed a dose-dependent inhibitory effect. Scale bar = 20 μm. **(B)** Quantification of NFAT1 nuclear entry, expressed as the percentage of cells exhibiting nuclear localization relative to the untreated, activated control cells. Data are presented as mean ± SD of independent biological replicates ****P<0.0001, ***P<0.001.

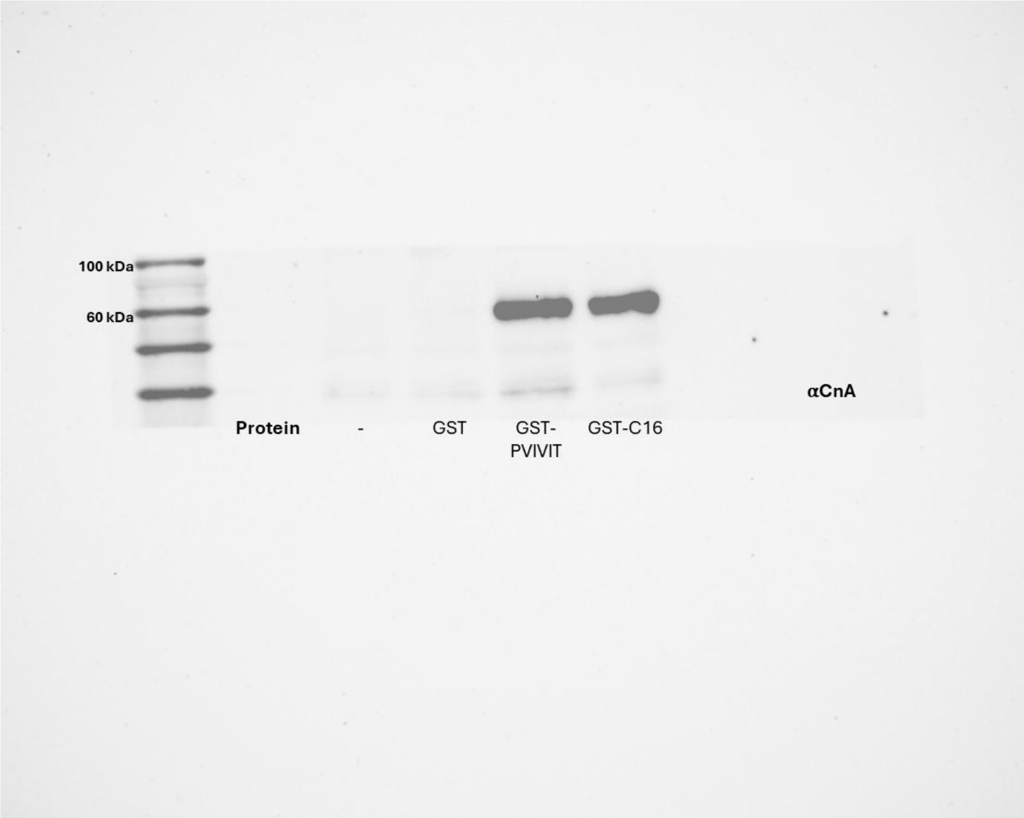


**Figure S3. Full blot of Fig. 2C top panel**

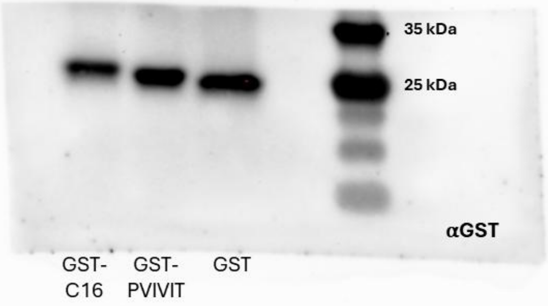


**Figure S4. Full blot of Fig. 2C middle panel**

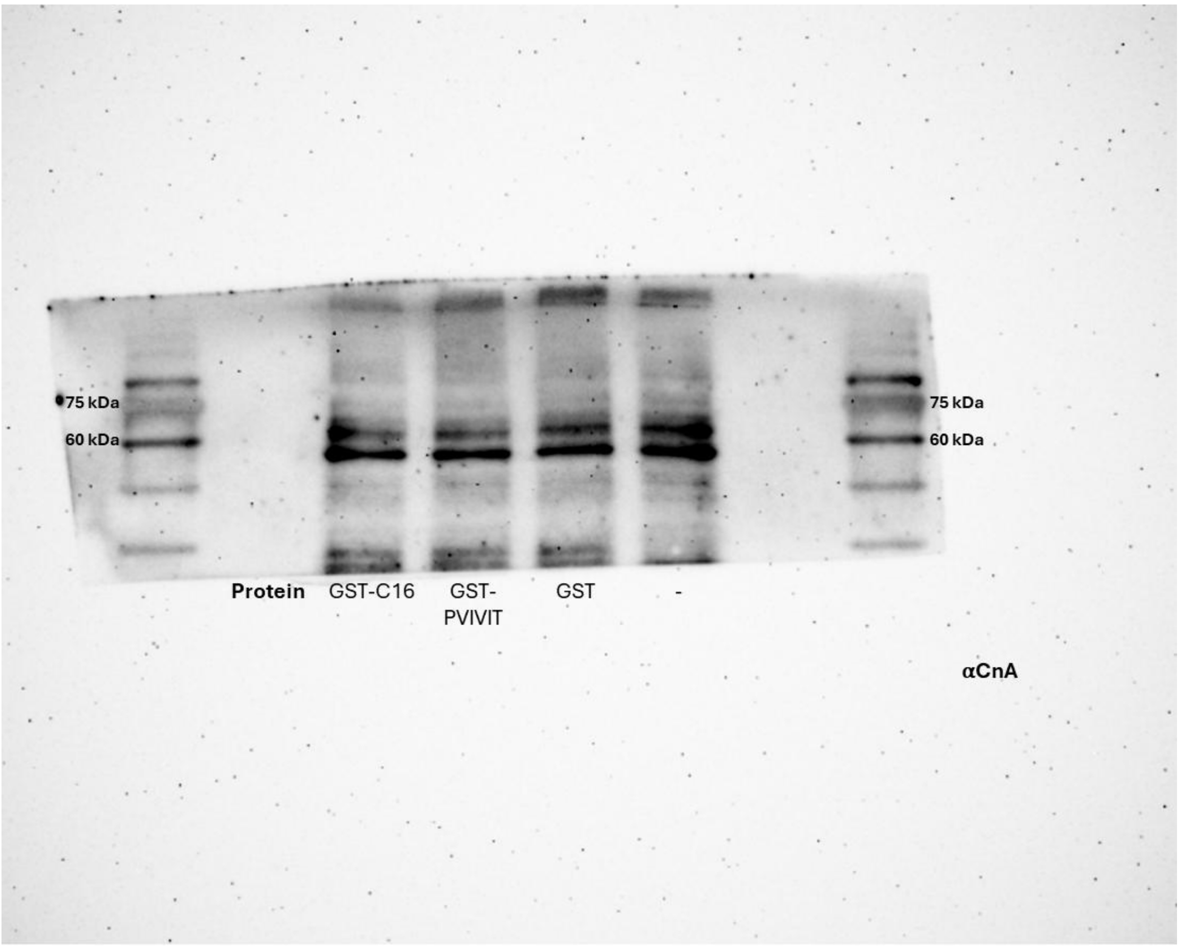


**Figure S5. Full blot of Fig. 2C bottom panel**
